# Surveying insect flower visitors to crops in New Zealand and Australia

**DOI:** 10.1101/373126

**Authors:** Brad G Howlett, Lisa J. Evans, Liam K Kendall, Romina Rader, Heather M. McBrydie, Samantha F.J. Read, Brian T. Cutting, Andrew Robson, David E. Pattemore, Bryony K. Willcox

## Abstract

The survey of insect flower visitors to crops dependent on their pollination is an essential component in determining their effectiveness as pollinators. In most cases, different survey techniques are required for different crops because of variation in planting design, floral density, spatial distribution of flowers or where additional factors such as the variation in plant vigour are being explored. Here we provide survey techniques that have been, or are currently being employed to survey flower visitors across different crops in New Zealand and Australia. Future studies may consider the use of similar designs that will allow for increased standardisation within and between locations and studies. This will provide opportunities for improved direct comparisons between studies, and the ability to combine data sets to address broader spatial-scale questions regarding insect pollination.

## Introduction

Understanding the abundance and identity of insect flower visitors is an important component in quantifying pollination within crops dependent on insect pollination. As part of research undertaken by The New Zealand Institute for Plant & Food Research Limited and the University of New England (New South Wales, Australia) to assess the role of managed honey bees and wild pollinators, we designed several standardised survey techniques to assess pollinator abundance and diversity within avocado (*Persea americana*), blueberry (*Vaccinium spp*.) kiwifruit (*Actinidia deliciosa*), macadamia (*Macadamia integrifolia* and *M*. *tetraphylla*) and mango (*Mangifera indica*) orchards located either in New Zealand or Australia. These add to the flower-visitor survey techniques for the insect seed crops onion (*Allium cepa*), carrot (*Daucus carota*), radish (*Raphinus sativus*), pak choi (*Brassica rapa* subsp. c*hinensis*) and white clover (*Trifolium repens*) that are described by Howett et al. (2009; 2013). Orchard design, the spatial distribution of flowers, and the need to design surveys to address the objectives of each study has required the development of multiple survey methods. These methods have been employed extensively across orchards to understand the abundance and diversity of insect flower visitors within and between orchard blocks and in some instances, between plants within blocks. Methods are described here in alphabetical order by crop type.

## 1. Avocados

Avocado flowers typically open twice, first as a pistillate (functionally female) flower and then as a staminate (functionally male) flower the following day. Staminate flowers can be distinguished by the positioning of the stamens with anthers (often dehiscing pollen) close to the pistil whereas, for pistillate flowers, stamens with non-dehiscing anthers are positioned prostrate against the petals. Flowers often open for relatively short periods during the day of time (commonly 2–4 hours) and this is dependent on prior weather conditions. Staminate and pistillate flowers within trees often overlap for a short period of time (usually not much longer than half an hour). Night flowering may also occur under cold overnight temperatures (near to or below 0°C).

Many commercial orchards consist predominantly or entirely of trees of a single cultivar. In New Zealand and southern Australia (New South Wales, Victoria and South Australia), the most common commercially grown cultivar is ‘Hass’. Additional polleniser cultivars may also be planted within the orchard blocks. These flower in the opposite phase to the fruiting trees, producing staminate flowers when the fruiting trees are producing pistillate flowers and vice versa. The aim of planting pollenisers is to assist pollination by producing pollen that can be transferred by pollinators to the fruiting trees. In many cases, fruit set is greater when the pollen is from a different cultivar to the female flower.

To help facilitate pollination, avocado growers often place hives of honey bees within their orchards, however, this is not universal. Other unmanaged flower visiting insect species have previously been noted visiting avocado flowers in New Zealand (Read et al. 2017) and Australia (Vithanage 1986; Evans et al. 2011). Our surveys methods described here include counts of insect flower visitor to both fruit and polleniser trees when present.

### Avocado survey method 1: New Zealand orchard blocks using a diagonal survey design

This survey method was designed to assess flower visitor abundances and diversity within and between orchard blocks across New Zealand with trees that could be highly variable flowering intensity during peak flowering time. The data may then be combined with measures of pollinator efficiency (e.g. Rader et al. 2009) to calculate pollinator effectiveness.

#### Flower-visitor surveys

The trees grown for commercial fruit production within the surveyed orchard blocks were of the cultivar ‘Hass’ but in most cases a small number of different and variable polleniser cultivars were also present. Blocks surveyed ranged in size from 0.6 ha to 3.4 ha.

#### Survey design

The survey design consisted of 18 ‘Hass’ trees per orchard, grouped into three sets of six trees. Within each set, three trees were located in a single row and the other three in a single adjacent row. Within each of these rows, every second tree was selected as a survey tree and survey trees were positioned diagonally across the two rows. Therefore, survey trees in row 1 were 1, 3, and 5 and in row 2 they were trees 2, 4 and 6 (Figure 1).

**Figure 1:**
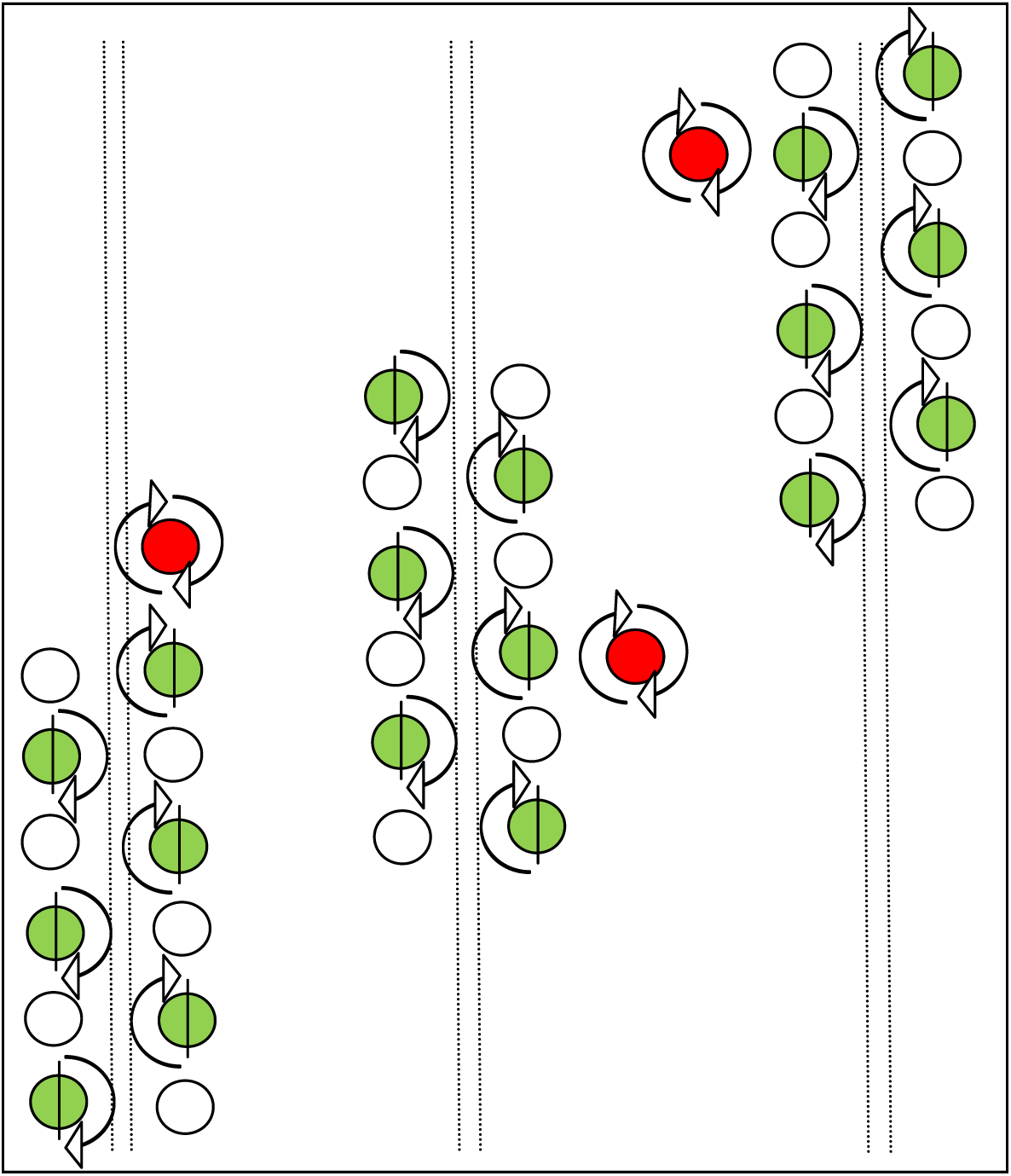
Survey design for assessing insect flower visitors in a New Zealand avocado orchard. Green circles represent the dominant commercial cultivar (‘Hass’), red circles pollenisers. Arrows represent 180°observation surveys on selected trees for ‘Hass’, 360°surveys on polleniser trees. Each survey was conducted on three sets of trees (two sets in opposing corners and one in the centre).

One tree set was located at a block corner, while a second set was located in the opposite corner. The third tree set was located in the centre of the block. The three sets were therefore positioned diagonally across the orchard block (Figure 1). When present, each polleniser tree selected for observation was located nearest to each ‘Hass’ tree set (Figure 1).

#### Survey times and flowering

For each daily survey, trees were observed three times during the day with surveys conducted from 0900–1000 h, 1200–1300 h, and 1500 and 1600 h, under fine weather conditions (15– 30°C) and when average wind speed was below 15 km/h. The flowering phase (staminate or pistillate flowers) of each tree was also recorded at each survey time. Wherever possible, surveys were conducted at a time of peak orchard flowering (mean staminate flower count > 40 staminate phase flowers/quadrat count on more than 50% of trees). To account for variation in flowering intensity between blocks and trees, surveys of flowers per unit area were also conducted within each observation tree (described in the section “Recording Flowering Intensity”).

#### Survey method

Only flowers between 0.5 m and 2.0 m above ground were surveyed for insect flower visitors. To restrict observations within this defined survey height, the observer vertically held a 1.5 m pole (dark green) near (25 ± 15 cm) the tree side. The observer then walked slowly around the tree observing inflorescences at a distance of within 30 cm, counting flower-visiting insects around half the trees circumference (180°) for a period of 120 ± 15 secs per tree. All flowers located from the outer canopy to the trunk of the tree were observed. This was possible because of the open nature of the canopy and the tendency for inflorescences to be located nearer the outer canopy of the tree than buried in foliage within the tree. Where a flower-visiting species was particularly abundant, a hand-held counter was used to record numbers. We surveyed the same sides of each tree located within the same row and the adjacent 180° of trees in the second row. Therefore, the side of each tree surveyed within each set faced inside the same row (Figure 1).

#### Recording flower visitors

Insects were recorded on a paper spreadsheet to species if possible. The paper spreadsheet was designed with listed taxa deemed most likely to be observed based on earlier preliminary observations. If an insect could not be identified through visual observation, the observer attempted to collect a specimen either directly into a jar containing tissue paper moistened with ethyl acetate for later identification or by first using a net from which the insect was transferred to a jar.

#### Measurement of weather variables

We measured wind speed, air temperature, relative humidity and light intensity (irradiance) (north, south and directly towards the sun). A Silva Windwatch was used to measure wind speed (km/h) over a 30 second period with minimum and maximum speeds recorded. Air temperature (°C) and relative humidity (%) were measured using a Thermo-Hydro recorder that was hung in full shade approximately 0.5 to 1.5 m above ground. Light intensity (watts/m^2^) was measured using a Daystar meter. All weather variables were measured just prior to conducting each insect flower-visitor survey (0900, 1200 and 1500h).

#### Recording Flowering Intensity

Flower-visitor surveys were coupled with surveys of flowering intensity that were conducted on the same day and on the same trees. Two separate counts for each tree were conducted during the day, one when trees were in their pistillate flowering phase and the other during their staminate phase. A single 75 × 75 cm^2^ quadrat was used to count open flowers within a segment of the area that was surveyed for insect flower visitors. The quadrat was placed at a height of 1.0 m above ground and positioned at the midpoint of the insect survey transect (i.e. facing directly into the row). A piece of tape attached to the quadrat side assisted in determining the correct height.

All inflorescences (budding, flowering, completed flowering) were initially counted separately within the area of the quadrat to the tree trunk when standing 50 cm directly behind the quadrat. Three inflorescences (regardless of the number of open flowers they contained) were then selected within the quadrat and marked with plastic tape. One in each of the following positions was tagged: (1) visually closest to the upper left-hand corner, (2) closest to the centre and (3) closest to the lower right-hand corner. To avoid sampling only inflorescences nearest to the quadrat (that was placed on the outer canopy), we chose the inflorescences that were closest to our quadrat points based on our field of view when standing 50 cm from the quadrat. Therefore, inflorescences close to the trunk could be selected if they were visually closest to the quadrat points. All open flowers were then counted on these inflorescences.

A count of inflorescences per quadrat and flowers (staminate and pistillate phase) per marked inflorescence could then be used to estimate of the number of flowers per area surveyed for each set of trees. This calculation could then be used to estimate numbers of flower visitors per flower.

*Recording other tree parameters*

The height (estimated) and radius (measured with a tape) of each tree was also noted. Where trees were irregular in shape, measures of the tree radius at its greatest and minimum extent were recorded.

### Avocado survey method 2: Australian orchard blocks using a diagonal survey design

This survey method was designed to assess flower visitor abundances and diversity within and between orchard blocks across orchards in South Australia, Victoria, New South Wales and Queensland. The method was often conducted simultaneously with avocado survey method 3. The design was modified from avocado survey method 1 to match method 3 more closely (observations conducted around entire tree circumferences). As with avocado survey method 1, the data may then be combined with measures of pollinator efficiency to calculate pollinator effectiveness.

#### Flower-visitor surveys

As in New Zealand, ‘Hass’ is the most commonly grown cultivar for commercial fruit production in South Australia, Victoria and New South Wales. In some of these orchards, a small number of different and variable polleniser cultivars are grown within blocks. Blocks surveyed ranged in size from 0.45 ha to 10.57 ha.

#### Survey design

The survey design consisted of nine ‘Hass’ trees per orchard, grouped into three sets of three trees. Within each set, two trees were located in the same row but separated by a single tree. The third tree was selected from an adjacent row but in a position adjacent to the unmarked centre tree separating the two survey trees. Therefore, survey trees in row 1 were 1 and 3 and in row 2, tree 2 (Figure 2).

**Figure 2:**
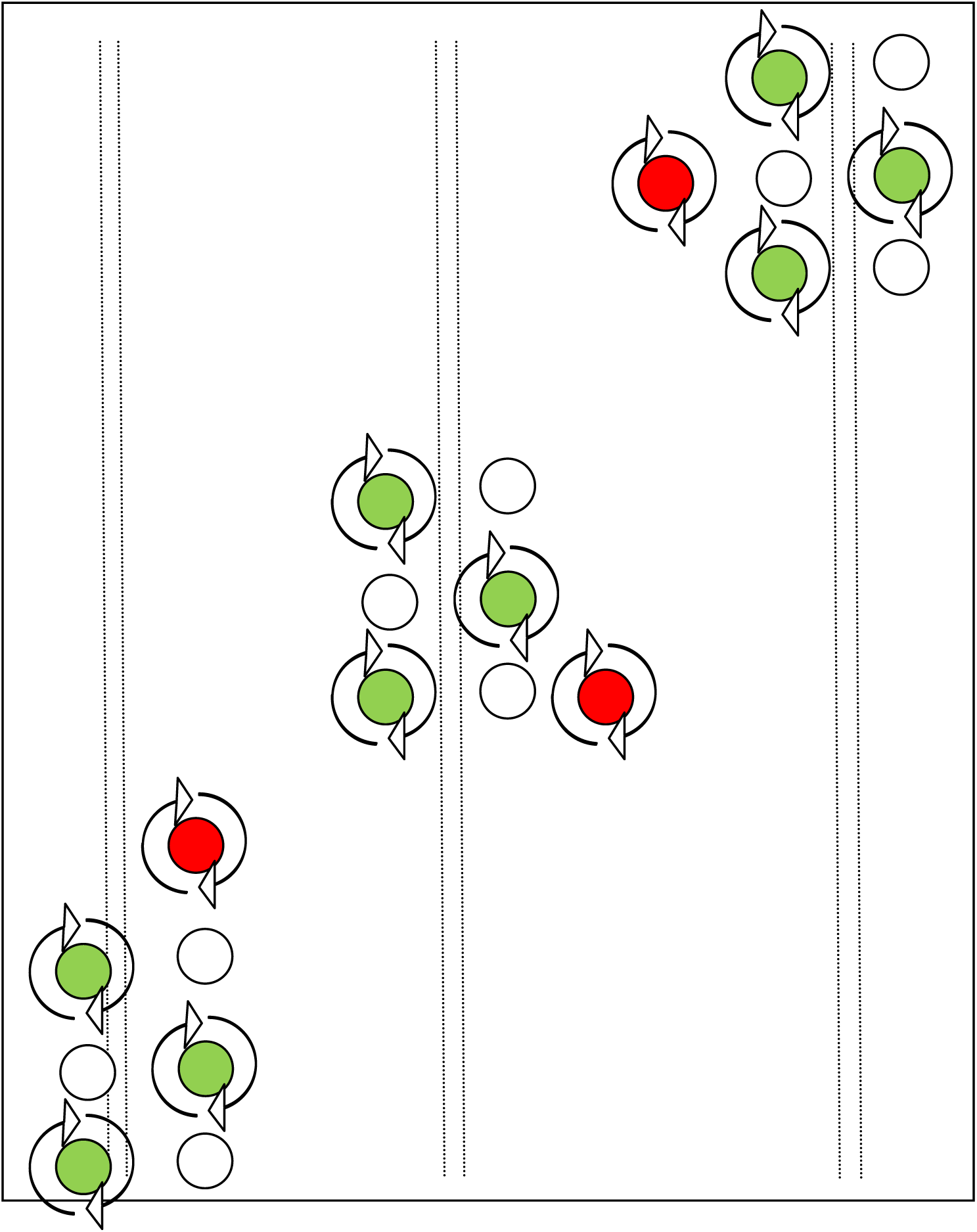
Survey design for assessing insect flower visitors in an Australian avocado orchard. Green circles represent the dominant commercial cultivar (‘Hass’), red circles pollenisers. Arrows represent 360°observation surveys on selected trees for ‘Hass’, 360° surveys on polleniser trees. Each survey was conducted on three sets of trees (two sets in opposing corners and one in the centre).

As for the New Zealand surveys, in each block, one tree set was located at a corner of a block, the second set in the opposite corner and the third set located in the centre of the block (Figure 2). In orchards with pollenisers, the polleniser tree nearest to each set of ‘Hass’ trees was selected for observation surveys (Figure 2).

#### Survey times and flowering

As in New Zealand, surveys were conducted three times per day (0900 –1000 h, 1200–1300 h, and 1500 and 1:00 h) under fine weather conditions (13–36°C) and when average wind speed was below 20 km/h. Flowering phase (staminate or pistillate flowers) of trees at survey time was also recorded. Wherever possible, surveys were conducted at peak flowering (mean staminate flower count > 40 staminate phase flowers/quadrat count on more than 50% of trees). To account for variation in flowering intensity between blocks and trees, surveys of flowers per unit area were also conducted within each observation tree (described in the section “Recording Flowering Intensity”).

#### Survey method

The survey method employed was mostly similar to that employed in New Zealand avocado orchards (avocado survey method 1). That is, flowers were surveyed for flower visitors between 0.5 m and 2.0 m above ground using a vertically held pole to limit survey height around the side of the tree. Observers surveyed flower visitors at a similar distance (within 30 cm) from inflorescences. However, an observer slowly walked around the full circumference (360°) of each tree (Figure 2) counting flower-visiting insects over a period of 240 ± 30 secs per tree.

As in New Zealand, the open canopy allowed all flowers located from the outer canopy to the trunk to be surveyed. A hand-held counter was used for very abundant flower-visiting taxa.

#### Recording Flower Visitors

The method used to record flower visitors was largely the same as described for avocado survey method 1. In summary, paper spreadsheets that were prepared following preliminary observations across four orchards to determine the more abundant flower visiting taxa.

However, in a small number of orchards blocks where flower-visiting species were very abundant, counts were recorded directly into an audio recorder. Unidentified insects were captured wherever possible as described for avocado survey method 1, to allow further identification.

#### Recording Flowering Intensity

As for New Zealand surveys, flowering-intensity surveys were conducted on the same day as flower-visitor surveys. Two surveys were conducted per day, one for each flowering phase (pistillate and staminate). The method was largely the same as employed in New Zealand blocks. That is, 75 cm^2^ quadrats were placed at a height of 1.0 m above ground and all inflorescences counted (flowering or not flowering), from outer canopy to trunk. Three inflorescences were chosen within the quadrat as described for avocado survey method 1.

Flower and inflorescence counts were conducted within quadrats placed on four sides of each tree. These were placed on the two opposite sides of the tree facing directly into the rows and the two sides facing the two neighbouring trees within the row. Thus, the four quadrat points were separated by 90°.

#### Recording other tree parameters

The height (estimated) and radius (measured with a tape) of each tree was also noted as described for avocado survey method 1.

#### Measurement of weather variables

Wind speed (minimum and maximum across a 30 second period), air temperature (°C), relative humidity (%) and light intensity (irradiance) W/m^2^ (north, south and directly towards the sun) were measured using the devices described for the avocado survey method 1.

### Avocado survey method 3: Australian orchard blocks using tree vigour classes

This survey method was designed to assess flower visitor abundances and diversity between specific trees categorised on the vigour of their growth.

#### Flower-visitor surveys

Surveys were conducted in orchards across two growing regions in Australia: Bundaberg, QLD 24.867° S, 152.351° E (within 100 km) and Renmark 34.174° S, 140.744° E (within 40 km), SA. Blocks within orchards that included only a single cultivar, in this instance ‘Hass’, were chosen for flower-visitor surveys. Blocks ranged in size from 7.3 ha to 32.38 ha in Bundaberg and 2.28 ha and 10.57 ha in Renmark.

#### Survey design

The survey design consisted of 18 trees per orchard. Trees were selected based on three classes of tree vigour (size and health) as defined by classified normalised difference vegetation index (NDVI) maps derived from a Worldview-3 satellite image (Robson et al. 2014). Six trees each, belonging to high-, medium- and low-vigour zones were identified from the classified NDVI maps before being verified by on-ground inspections. Trees of each class were chosen to ensure even representation throughout the block. Trees were then flagged using plastic tape.

#### Survey times and flowering

Surveys were conducted three times per day (0900 –1000 h, 1200–13:00 h, and 1500 and 1600 h). Flowering phase (staminate or pistillate flowers) of trees at survey time were also recorded.

#### Survey method

The method used was the same as for avocado survey method 1. In summary, flowers were surveyed for flower visitors at a height of between 0.5 m and 2.0 m above ground. The observer slowly walked around the full circumference (360°) of each tree observing flower visitors at a distance within 30 cm of inflorescences. The time spent to complete one tree survey was 240 ± 30 secs.

The open canopy allowed all flowers located from the outer canopy to the trunk to be surveyed. A hand-held counter was used for very abundant flower-visiting taxa.

#### Recording Flower Visitors

Similar to avocado survey methods 1 and 2, insects flower visitors in orchard blocks were recorded on paper spreadsheets prepared following preliminary observations across a small number of orchard blocks. Unidentified insects were captured where possible to allow for later identification.

#### Recording Flowering Intensity

Flowering-intensity surveys were the same as described for the avocado survey methods 1 and 2. That is, they were conducted on the same day as flower-visitor surveys. Two surveys were conducted per day, one for each flowering phase (pistillate and staminate). The method consisted of 0.75 m^2^ quadrats being placed at a height of 1.0 m above ground and all inflorescences counted (flowering or not flowering) from outer canopy to trunk. Three inflorescences were chosen within the quadrat as described for method 1.

Flower and inflorescence counts were conducted within quadrats placed on four sides of each tree. These were placed on the two opposite sides of the tree facing directly into the rows and the two sides facing the two neighbouring trees within the row. Thus, the four quadrat points were separated by 90°.

#### Measurement of weather variables

Wind speed (minimum and maximum across a 30 second period), air temperature (°C), relative humidity (%) and light intensity (irradiance) W/m^2^ (north, south and directly towards the sun) were measured using the devices described for method 1.

### Avocado survey method 4: Australian orchard blocks using single transect

#### Flower-visitor surveys

Flower-visitor surveys were conducted in the same Bundaberg ‘Hass’ orchard blocks that were used the previous year for avocado (avocado survey method 3). The method was altered to increase the numbers of flowers surveyed, thereby, increasing the chance of recording higher flower visitor counts and gain further knowledge of the diversity of the flower visitor assemblage.

#### Survey design

The survey design consisted of a 50 m transect per orchard block. Transects were selected away from the edges of orchards to reduce potential edge effects on pollinator abundances and yield. Trees at the start and end of each transect were then flagged using plastic tape.

#### Survey times and flowering

Surveys were conducted four times per day (0800 –0820 h, 1000–1020 h, 1200–1220 h and 1600–1720 h). The flowering phase (staminate or pistillate flowers) of each tree at each survey time was also recorded.

#### Survey method

The survey method employed consisted of walking the 50 m transect for a duration of 20 mins at each of the designated survey times. An observational scan (180°) of each tree (similar to that shown in Figure 1) within a transect was conducted for flower visitors, with the observer also scanning to the height of the canopy from the ground.

Flowers located from the outer canopy to the trunk were surveyed. A hand-held counter was used for very abundant flower-visiting taxa.

#### Recording Flower Visitors

Insects flower visitors in most orchard blocks were recorded on paper spreadsheets to species level where possible. For insect species that could not be identified by the observer, a collection of the specimen was attempted using the same methods described for avocado survey method 1.

#### Recording Flowering Intensity

Flowering-intensity surveys were conducted on the same day as insect surveys using similar methods described for avocado survey method 1. That is, two surveys were conducted per day, one for each flowering phase (pistillate and staminate) using 0.75 m^2^ quadrats

Flower and inflorescence counts were conducted within quadrats placed on two sides of each tree. These were placed on the two opposite sides of the tree facing directly into the rows. Thus, the two quadrat points were separated by 180°.

#### Measurement of weather variables

Air temperature (°C) and relative humidity (%) were measured at the beginning and end of each survey using the devices described for avocado in survey method 1.

## 2. Blueberry

### Blueberry Survey Method 1: Australian blocks using a transect survey

Blueberry flowers are urceolate in shape, small in size (corolla length = 11mm and aperture = 3.5mm (Lyrene 1994)) and hermaphroditic. They typically remain open for about three days. In open flowers, the anthers remain enclosed within the corolla whereas the stigma protrudes slightly above the corolla aperture.

Commercial blueberry growers in New South Wales typically grow two blueberry species: Southern Highbush (*Vaccinium corymbosum*) and Rabbiteye (*V*. *ashei* or *virgatum*). Flowering phenology differs between Southern Highbush and Rabbiteye: Southern Highbush peak flowering occurs from April to July. In Rabbiteye, it occurs from late August to early October. Varietal planting patterns differ for Southern Highbush and Rabbiteye. Southern Highbush varieties are grown in a mosaic of single cultivar one ha^-1^ blocks whereas Rabbiteye blocks commonly consist of two or three varieties planted in successive rows.

Very little is known about the blueberry pollinator community in Australia. Honeybees are viewed as the primary pollinators (e.g. Goodman & Clayton-Green 1988) and are stocked at 4-7 hives ha-1 to ensure adequate fruit set. Preliminary results show that stingless bees (*Tetragonula carbonaria*) and hoverflies also visit flowers.

*Flower visitor surveys*

Surveys were conducted in blocks of both Southern Highbush and Rabbiteye blueberries during peak flowering periods in 2016 and 2017. They were conducted in blocks of two blueberry species: Southern Highbush (*Vaccinium corymbosum*) and Rabbiteye (*V*. *ashei* or *virgatum*). Southern Highbush was surveyed in May 2016 and May 2017 and RE in September 2016 and September 2017. Surveys were conducted primarily on the North Coast, NSW, on farms located between Woolgoolga (30.112259°S, 153.190375°E) and Dirty Creek (29.990232°S, 153.143171°E). Two farms in Atherton, Far North Queensland (17.243906°S, 145.478321°E), each growing Southern Highbush blueberries, were also surveyed in 2017.

#### Survey design and timing

Surveys were conducted across a 50m transect along one crop row in each block. Plant spacing varied between 0.9 – 1m, with between 50 −55 plants being surveyed across each transect. We selected the transect row at random, using a random number generator, excluding the first and last five rows.

#### Survey times and flowering

Surveys were conducted over three periods during each day: 1000 - 1020 h, 1200 - 1220 h and 1400 - 1420 h and each block was surveyed over three days.

#### Survey method

Blueberry plants in cultivation are typically between 1.5 – 2.5m in height. The entirety of the ‘row-side’ of each plant was scanned for flower visitors. Given the plant spacing, a maximum of 20-25 seconds was spent observing floral visitors on each plant depending on floral abundance.

#### Recording Flower-Visitors

As well as recording flower visitors during each survey, we also recorded foraging behaviour (i.e. nectar or pollen foraging). We denoted behaviour based on the presence (or absence) of corbiculae (pollen baskets), in the case of honeybees and stingless bees, or pollen on the underside of the thorax and abdomen. Behaviour was not recorded for hoverflies. Following each survey, visitors were collected by hand. We also obtained honeybee stocking rates from each landowner

#### Recording Flowering Intensity

Counts of flowering intensity were conducted once within each block surveyed. We selected five rows and five plants within each row at random, using a random number generator. On each plant, we then recorded the total number of open flowers.

#### Measurement of weather variables

Weather variables: Daily readings of air temperature (°C), relative humidity (%), wind speed (km/h), light intensity and precipitation (mm) data were collected from the weather stations located within the main Costa Berry Farm complex in Dirty Creek, NSW and Atherton, Queensland.

## 3. Kiwifruit

Kiwifruit produce staminate or pistillate flowers contained on separate vines. Pistillate flowering vines (fruiting vines) of a specific cultivar are planted within an orchard block but often a smaller number of staminate vines of a different cultivar are interspaced throughout a block. Wire trellising is used to support vines and the most common trellis design employed in older orchards is the T-bar system, although newer orchards often use a pergola design (New Zealand Kiwifruit Growers Incorporated 2016). Inflorescences typically contain up to three flowers. In T-bar trellised orchards, flowers usually face downward (towards the ground), at a height of 1.7 to 2 m above ground. Pistillate flowers contain around 40 styles surrounded by numerous anthers producing sterile pollen. Staminate flowers do not contain styles but have anthers that produce fertile pollen. Historically ‘Hayward’ (*Actinidia chinensis* var. *deliciosa*) was the most commonly grown cultivar and in 2016 was still the cultivar producing the largest number of fruit by volume in New Zealand (New Zealand Kiwifruit Growers Incorporated 2016). Although wind is considered a vector for pollination, insects, particularly honey bees, are regarded as essential for maximising pollination within New Zealand kiwifruit orchards. Hives of honey bees are regularly placed within orchards. In addition to honey bees, other bee species, flies and beetles visit kiwifruit flowers in New Zealand (Howlett et al. 2017).

### Kiwifruit survey method 1: New Zealand orchard blocks using a diagonal survey design

#### Flower-visitor surveys

Our study was conducted in fully mature, flowering ‘Hayward’ orchard blocks containing evenly spaced staminate polleniser vines. Orchards were located in three regions, within a 100 km radius of Katikati (37°33’S 175°55’E), within 50 km of Gisborne (38° 40’S, 178° 01’E) and within 50 km of Hastings (39° 39’ S, 176° 50’ E).

#### Survey design

All blocks surveyed had been planted using the T-bar trellis system. Flowers on ‘Hayward’ pistillate vines were assessed by walking four sets of transects underneath the vines positioned throughout each block, while flowers from four staminate vines (various cultivars) nearest to the pistillate vine transects were also assessed.

The survey design within each block consisted of four within row transects, spaced diagonally and equidistantly from one corner to the opposing corner (Figure 3). The two opposing corner transects were located in the first row at the point where the first wire trellis commenced. The two inner transects were spaced equidistantly between the corner transects.

**Figure 3:**
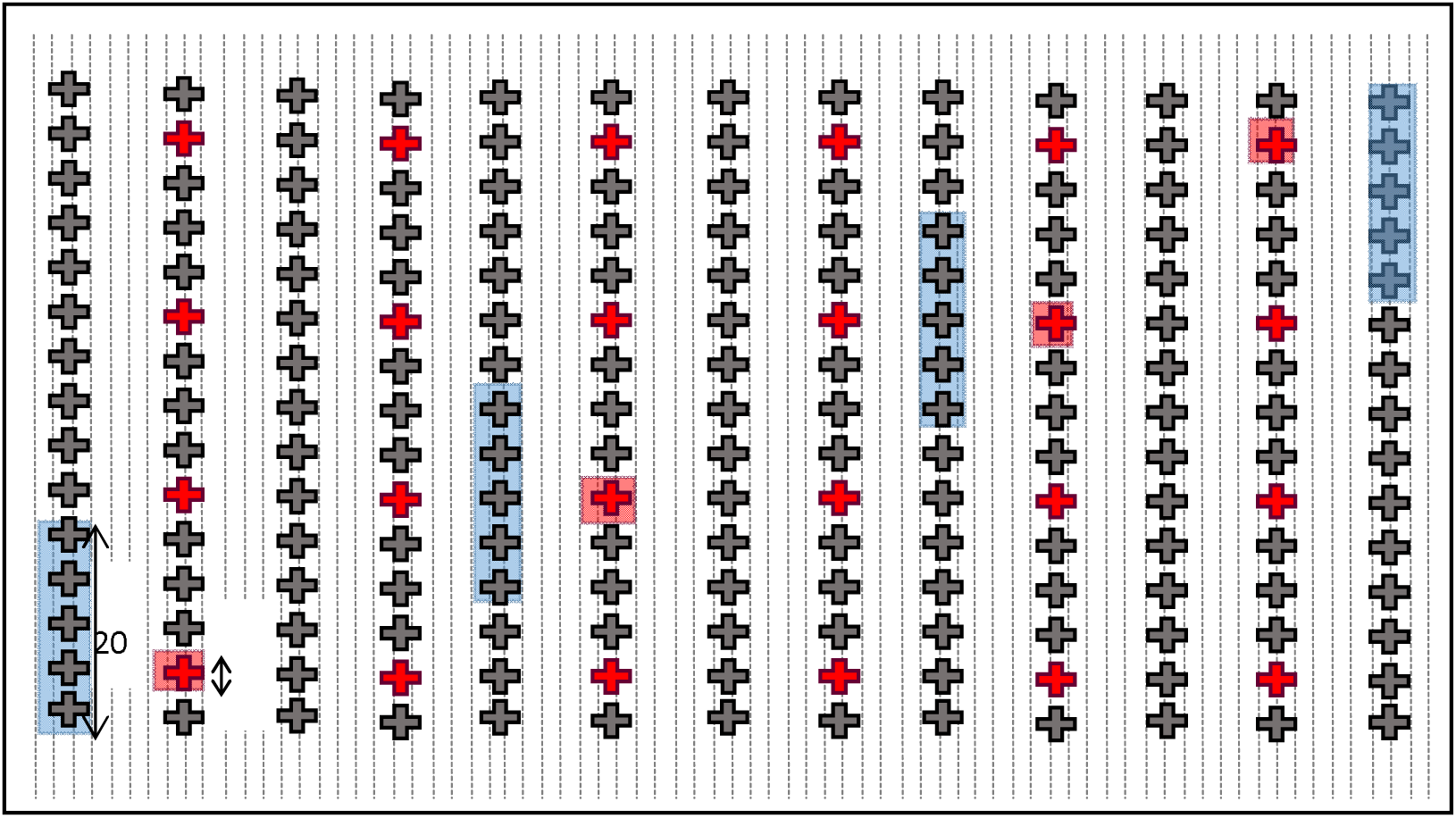
Survey design for assessing insect flower visitors in a New Zealand kiwifruit orchard. Grey crosses represent the dominant commercial cultivar (‘Hayward’ Actinidia chinensis var. deliciosa), red crosses, pollenisers. Blue shading represent observational transects on ‘Hayward’ (length 20 m width 1.3-1.4 m), red shading, are transects on polleniser vines (3.6 x 0.4 m).

Each ‘Hayward’ transect was 20 m in length and between 1.2 m and 1.4 m in width (determined by the distance between supporting trellis wires). The actual area of each transect was determined by measuring the width of the transect at the start and at the end of the transect length. If the measurement varied between the measurements (never more than 5 cm difference recorded), the mean between the two was used to calculate transect area.

Single staminate vines were spaced evenly throughout orchard blocks surveyed and the nearest flowering vine to each ‘Hayward’ vine transect was chosen to survey (Figure 3). The survey length for each vine was 3.6 m (1.8 m on each side of the vine trunk) and the width was 0.4 m. Lengths and widths were measured and marked with tape.

#### Survey times and flowering

Each block was surveyed for one hour at three times throughout the day from 1000–1100 h, 1200–1300 h and 1400–1500 h under fine weather conditions (15–30°C) and when average wind speed was below 15 km/h. Where possible, orchard block surveys were conducted at peak flowering (density of fully open flowers > 7/m^2^ in ‘Hayward’ transects). To account for variation in flowering intensity between blocks, surveys of flowers per unit area were also conducted within each transect (see section “Recording Flowering Intensity” below).

#### Survey method

Insect flower visitors to fully open flowers (excluding buds and flowers that had lost all petals) were observed from within a distance of 30 cm of each flower by the observer walking slowly beneath the canopy. The time spent observing flower visitors along each transect was 12 ± 2 mins for ‘Hayward’ transects and 4 ± 1 mins for staminate vines.

#### Recording Flower Visitors

Insects flower visitors in orchard blocks were recorded on paper spreadsheets as outlined for avocado survey method 1. Where possible, unidentified specimens were collected, again using the same methods employed in avocado method 1.

#### Recording Flowering Intensity

Flowering-intensity surveys were conducted on both ‘Hayward’ and polleniser flowering vines on the same day that each insect survey was completed. These were conducted inside the area of each insect survey transect.

Flower counts were conducted within three square quadrats (area 0.81 m^2^) within each ‘Hayward’ transect. Quadrats were positioned at 5, 10 and 15 m along the transect length and equidistant between the transect width. All fully open flowers (containing petals), old flowers (no petals) and buds (anthers/stamens not visually exposed) were counted within each transect.

For staminate vines, flower counts were conducted within two rectangular quadrats of area of 0.36 m^2^ (0.9 m length × 0.4 m wide). Staminate vines typically contained two dominant branches pruned to run along each side of the trellis. Each transect was positioned on either side of the vine trunk to incorporate flowers between 0.9 m to 1.8 m from the trunk. As with ‘Hayward’ vines, fully open flowers, old flowers and buds were counted within the transect area.

#### Measurement of weather variables

Wind speed, air temperature, relative humidity and light intensity were measured using the devices described for the avocado survey method 1.

## 4. Macadamia

Queensland and New South Wales are the largest producers of macadamia nuts in Australia. Macadamia flowers are hermaphroditic (releasing viable pollen before the stigma becomes receptive) and also produce nectar. Prior to flower opening, pollen is dehisced from the anthers onto a bulbous region located at the tip of the style known as the pollen presenter. The stigma is located at the tip of the pollen. Macadamia produce approximately 100—300 florets within a raceme and several thousand racemes per tree during a flowering season. There are numerous cultivars of macadamia and these are grown as single cultivar or multiple cultivar blocks. In multiple cultivar blocks, growers commonly grow alternating rows of a different cultivar but growers may also grow different cultivars within the same row. Our surveys were conducted mostly within single cultivar blocks. Honey bee and stingless bee hives are occasionally placed within orchards, however, the numbers of hives growers have placed within their blocks can vary greatly. In Australia Honey bees and stingless bees are considered the most abundant pollinators but many other species, particularly beetles and flies also visit flowers (Vithanage & Ironside 1986; Heard & Exley 1994) The review articles by Trueman (2013) and Howlett et al (2015) provide further detail on macadamia floral biology and pollination.

### Macadamia survey method 1: Australian orchard blocks using a diagonal survey design

The survey method was designed to assess flower visitor abundances and diversity within and between orchard blocks in Queensland and New South Wales (Australia).

#### Flower-visitor surveys

Surveys were conducted in flowering orchard blocks consisting of one cultivar, with the exception of two blocks that contained two cultivars. Cultivars surveyed were ‘741’ (19 blocks), ‘344’ (12 blocks), ‘842’ (eight blocks), ‘816’ (one block), and ‘Daddow’ (five blocks). The mixed cultivar blocks contained the cultivars ‘344’and ‘842’ in block 1 and ‘741’and ‘344’ in block 2. Surveys were conducted from 31 August to 19 September 2015, across three different regions in Australia. From north to south: seven orchards near Bundaberg (within 100 km of Bundaberg, 24.8°S 152.3°E), 10 orchards in Gympie – Glasshouse (within 200 km of Gympie – Glasshouse Mountains, 26.2°S 152.6°E– 26.9°S 152.9°E), and six orchards in the Northern Rivers of New South Wales (within 100 km of Lismore 28.8°S 153.4°E).

#### Survey design

Twelve trees were observed per orchard in four sets of three trees. Each set of trees consisted of three neighbouring trees in a single row. The first set of three trees were located in the corner of the block (row one), trees 1, 2, and 3, while another set of three in the opposite corner (final row), last three trees (Figure 4). The other two sets of trees were located equidistant within the orchard to form a diagonal survey of three grouped trees across the orchard (Figure 4).

**Figure 4:**
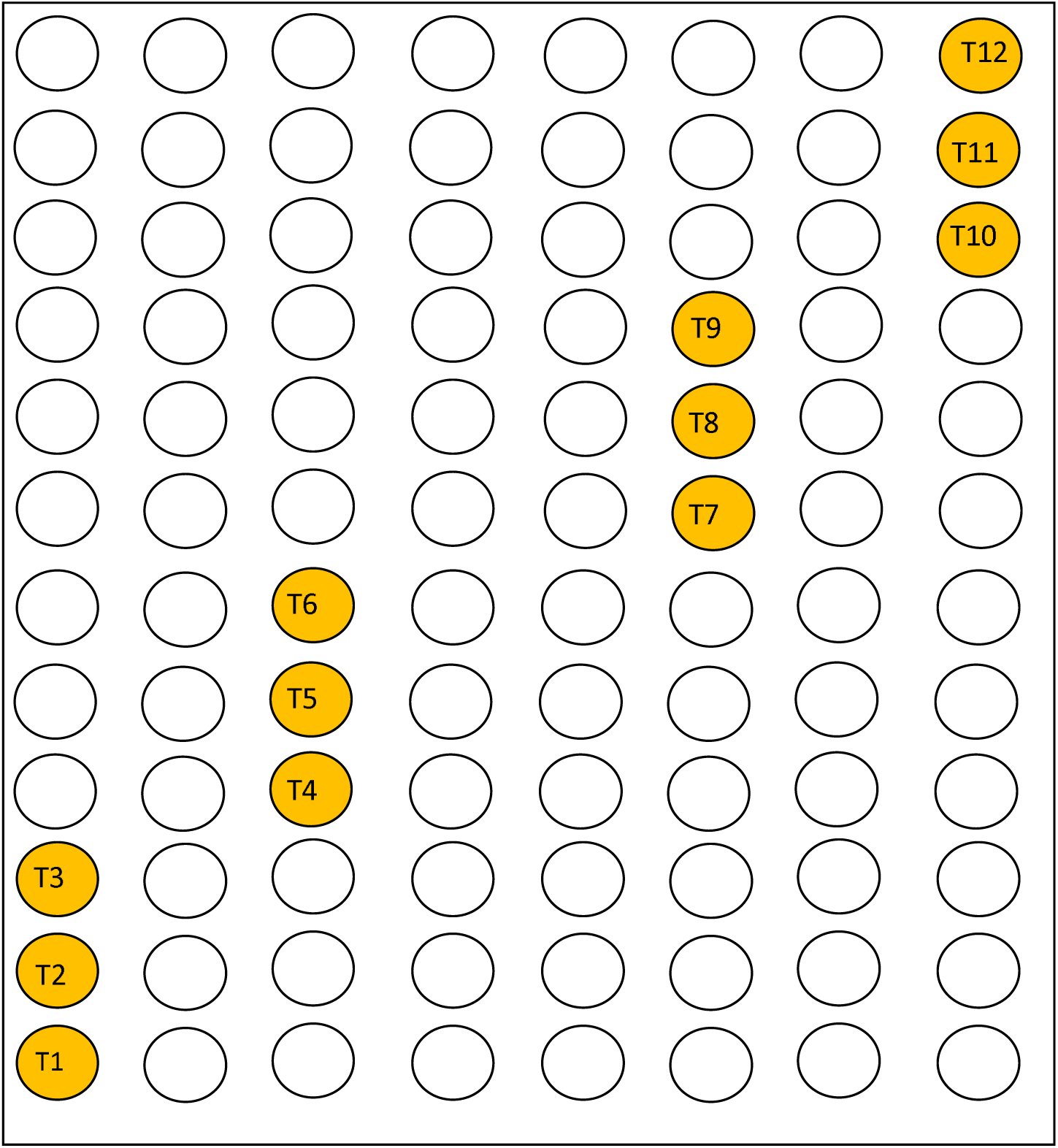
Survey design for assessing insect flower visitors in an Australian macadamia orchard. Orange circles with numbers represent surveyed trees throughout the block.

#### Survey times and flowering

For each survey, trees were observed three times during the day from 0830–0930 h, 1200– 1300 h and 1500–1600 h. Surveys were conducted under fine weather conditions (17–33°C) and when average wind speed was below 24 km/h. Where possible, orchards surveys were conducted when the block was considered at peak flowering (more than 50% of tree with > 4 flowering racemes/m^2^). To account for variation in flowering intensity between blocks and trees, surveys of flowering racemes were also conducted within each observation tree (see recording flowering intensity below).

#### Survey method

The survey method was restricted to the side of each tree as the canopy was unable to be accessed from the ground. During each survey, a 1.5 m long pole (as described in avocado survey method 1) was held with the base at a vertical height of 0.5 m above the ground (as determined by string attached to the pole base). The pole was also equipped with a perpendicular horizontal wooden pole of length 0.75 m. This was used as a guide to restrict insect counts on racemes to the area between the outer canopy and 0.75 m within the canopy. This was necessary as racemes could be hidden deeper within the foliage of the canopy and difficult to observe directly. In some cases where the base of the foliage had been pruned approximately 1 m above the ground, it was necessary to hold the pole base at a higher point in the foliage to ensure the area containing flowering racemes remained equivalent to other orchards. The observer then slowly walked around the entire circumference of each tree counting flower-visiting insects at a distance within 30 cm. Care was taken to avoid the pole brushing the foliage that potentially could disturb flower visitors. Where a particular flower-visitor was particularly abundant, a hand-held counter was used to count numbers.

#### Recording flower visitors

Flower visitors were recorded on a paper spreadsheet to species if possible. The spreadsheet was designed to contain specific taxa deemed most likely to be observed based on earlier preliminary observations as well as knowledge obtained from previous studies by Vithanage and Ironside (1986) and Heard and Exley (1994). Wherever possible, collections were made of flower-visiting insects that could not be identified using the same methods described for avocado survey method 1

#### Recording Flowering Intensity

To assist in evaluating flowering intensity, counts of flowering racemes (inflorescence) on each survey tree were also conducted during the day of survey. Counts were conducted on four sides of each tree, two facing directly into each adjacent row and two along the tree row on opposite sides of the tree. Thus, the quadrats were placed on the side of each tree 90° apart. Quadrats were designed to measure the number of racemes contained within a cubic volume of 0.75m^3^. The design consisted of two square quadrats (constructed using wooden stakes) that were attached together along one side. These could be folded outwards to assist in defining the area within which racemes were then counted. The quadrat was held at a height of 1.5 m (inside the transect area of the flower-visitor surveys), and separate counts were conducted for budding racemes, racemes with less than 10 open flowers, fully flowering racemes and racemes that had completed flowering.

#### Recording other tree parameters

The height (estimated) and radius (measured with a tape) of each tree was also noted as described for avocados method 1. Photographs of each tree were also taken from opposing sides from within the tree rows.

#### Measurement of weather variables

Wind speed, air temperature, relative humidity and light intensity were measured using the devices as described for avocados method 1.

### Macadamia survey method 2: Australian orchard blocks using tree vigour classes

#### Flower-visitor surveys

Blocks surveyed within orchards contained a single cultivar ‘741’. The blocks ranged in size from 5.22 ha to 20.79 ha.

*Survey design*

The same survey design as used for avocado survey method 3 was employed. That is, the survey design consisted of 18 trees per orchard. Trees were selected based on three classes of tree vigour (size and health) as described for avocado survey method 3. Six trees from each of the vigour zones, high, medium and low, were identified from the classified NDVI maps before being verified by on ground inspections. Trees of each class were chosen to ensure even representation throughout the block. Trees were then flagged using plastic tape.

#### Survey times and flowering

The methods used were the same as macadamia survey method 1. Surveys were conducted three times per day (0900–1000 h, 1200–1300 h, and 1500–1600 h).

#### Survey method

Methods were similar to those described for macadamia survey method 1. That is, flowering racemes from the canopy edge to a depth of 0.75 m within the canopy and at a height of between 0.5 and 2.0 m were surveyed around the full circumference of each tree. The observer surveyed flower visitors from within 30 cm of flowering racemes and spent 180 ± 30 secs observing each tree.

#### Recording Flower Visitors

Methods were the same as those used for macadamia survey method 1. That is, insect flower visitors were recorded on paper spreadsheets that included insect flower visiting taxa which commonly visit macadamia racemes. Unidentified flower visitors were captured where possible for later identification.

#### Recording Flowering Intensity

Flowering-intensity surveys methods were the same as those used for macadamia survey method 1. That is, the number of racemes (budding, early flowering, fully flowering and finished flowering) were counted within a canopy volume of 0.75 m^3^ on each of four sides (separated by 90°) of the canopy, 1.0 m above ground level.

#### Measurement of weather variables

Wind speed (minimum and maximum across a 30 second period), air temperature (°C), relative humidity (%) and light intensity (irradiance in W/m^2^) (north, south and directly towards the sun) were measured using the devices described for the New Zealand avocado surveys.

### Macadamia survey method 3: Australian orchard blocks using single transect survey

#### Flower-visitor surveys

Flower-visitor surveys were conducted in the same orchard blocks (cultivar ‘741’) that were used in the previous year for the tree vigour surveys (Macadamia survey method 2).

#### Survey design

The survey design consisted of a 50 m transect per orchard block. Transects were selected a minimum distance of 25 m away from the edges of orchards in order to reduce the potential effect of the edge on flower visitor abundance and diversity. Trees at the start and end of each transect were then flagged using plastic tape.

#### Survey times and flowering

Surveys were conducted four times per day (8:00–8:20, 10:00–10:20, 12:00–12:20, and 16:00–16:20 hr).

#### Survey method

The survey method employed for all three crops consisted of walking the 50 m transect for a duration of 20 min at each of the designated survey times. An observational scan (180°) of each tree (similar to that shown in Figure 1) within a transect was conducted for flower visitors, with the observer scanning to the height of the canopy from the ground.

Flowers located from the outer canopy to the trunk were surveyed. A hand-held counter was used to count very abundant flower-visiting taxa.

#### Recording Flower Visitors

Insects flower visitors in most orchard blocks were recorded and collected using the same method described in avocado survey method 1.

#### Recording Flowering Intensity

Flowering-intensity surveys were conducted on the same day as flower-visitor surveys. One survey was conducted per day. As with macadamia survey method 1, all racemes (buds, less than 10 flowers open, fully flowering or finished flowering) were counted within a volume of a 0.75 m^3^ using fold-out quadrats placed at a height of between 1m–1.5 m above the bottom of the canopy.

Raceme counts were conducted within quadrats placed on two sides of each tree. These were placed on the two opposite sides of the tree facing directly into the rows. Thus, the two quadrat points were separated by 180°.

#### Measurement of weather variables

Air temperature (°C) and relative humidity (%) were measured at the beginning and end of each survey using the devices described for avocado survey method 1.

## 5. Mango

Mango is predominantly grown commercially throughout northern Australia (Queensland and Northern Territory) but are also grown in New South Wales, Western Australia and Victoria. Flowers are produced on panicles often containing several hundred to several thousand flowers. The majority of flowers produced by mango trees are staminate while the remainder are hermaphroditic. Self-pollination and cross-pollination may result in fruit set, however, insect pollination is considered important for maximising yields in many cultivars. In Australia, wasps, bees, ants and flies are frequent flower visitors and have been considered pollinators (Anderson et al. 1982). The review by Ramirez & Davenport (2016) provides further detail of mango floral morphology, reproductive biology and pollinators.

### Mango Survey Method 1: Australian orchard blocks using single transect survey

#### Flower-visitor surveys

Surveys were conducted in mango orchards that contained one of the cultivars ‘Calypso’ (Bundaberg, QLD) and ‘Keitt’ (Mareeba, QLD). Mango orchard blocks in Bundaberg ranged in size from 1.76ha–3.77ha. In Mareeba, mango surveys were conducted in orchards that grew multiple cultivars. While some had blocks of ‘Keitt’ as the only cultivar, many grew the cultivar in single rows or in rows that alternated with other cultivars. In these orchards, we restricted surveys accordingly. That is, within single rows of ‘Kiett’ or on ‘Kiett’ trees within rows containing mixed cultivars.

#### Survey design

The survey design consisted of a 50 m transect per orchard block or orchard. Transects were selected away from the edges of orchards in order to reduce edge effects. For blocks containing other cultivars within the row, we avoided the non ‘Kiett’ trees and recommenced the transect at the next ‘Kiett’ tree. In this case the transect length was extended so that 50 m of only ‘Kiett’ trees were observed. Trees at the start and end of each transect were then flagged using plastic tape.

#### Survey times and flowering

Surveys were conducted four times per day (0800–0820 h, 1000–1020 h, 1200–1220 h, and 1600–1620 h).

#### Survey method

The survey method employed consisted of walking the 50m transect for a duration of 20 mins at each of the designated survey times. An observational scan (180°) of each tree (similar to that shown in Figure 1) within a transect was conducted for flower visitors, with the observer scanning the height of the canopy from the ground.

Flowers located from the outer canopy to the trunk were surveyed. A hand-held counter was used for very abundant flower-visiting taxa.

#### Recording Flower Visitors

Flower visitors were recorded and where necessary, collected, using the methods described for avocado survey method 1.

#### Recording Flowering Intensity

Flowering-intensity surveys were conducted once for each orchard coinciding with a pollinator survey day. The method consisted of recording the number of primary branches emanating the trunk of each tree, then the number of open and closed panicles (inflorescence) on each of three primary branches and the number of hermaphroditic and staminate flowers on 10 randomly chosen panicles per tree. This was done for each tree located within the orchard transect.

#### Measurement of weather variables

Air temperature (°C) and relative humidity (%) were measured at the beginning and end of each survey using the devices described for avocado survey method 1.

## Acknowledgements

This work has been funded and supported by:

- Horticulture Innovation Australia through the projects MT13060 ‘Optimising pollination of macadamia & avocado in Australia’ and PH15000 ‘Strengthening and Enabling Effective Pollination for Australia’ with co-investment from Plant and Food Research, Australian Macadamia Society and funds from the Australian Government.
- The University of New England and funds from RnD4Profit-14-01-008 “Multi-scale monitoring tools for managing Australian Tree Crops: Industry meets innovation”.
- Ministry of Business, Innovation and Employment project C11X1309 ‘From Bee minus to Bee Plus and Beyond’ with co-funding from Zespri Kiwifruit, New Zealand Avocados, Foundation for Arable Research and Heinz Watties.

We also thank Avocados Australia, the Australian Mango Industry Association, Costa group, and the many growers and support staff that together, have allowed us to commence using these methods across the many crops and blocks.

